# Age-related reductions in the number of serial sarcomeres contribute to shorter fascicle lengths but not elevated passive tension

**DOI:** 10.1101/2020.12.21.423814

**Authors:** Geoffrey A. Power, Sean Crooks, Jared R. Fletcher, Brian R. Macintosh, Walter Herzog

## Abstract

We investigated age-related changes to fascicle length (F_L_), sarcomere length (SL), and serial sarcomere number (SSN), and how this affects passive force. Following mechanical testing to determine passive force, the medial gastrocnemius muscle of young (*n=*9) and old (*n=*8) Fisher 344BN hybrid rats was chemically fixed at the optimal muscle length for force production; individual fascicles were dissected for length measurement, and laser diffraction was used to assess SL. Old rats had ∼14% shorter F_L_ than young, which was driven by a ∼10% reduction in SSN, with no difference in SL (∼4%). Passive force was greater in the old compared to young rats at long muscle lengths. Shorter F_L_ and reduced SSN in the old rats could not entirely explain increased passive forces for absolute length changes, owing to a slight reduction in SL in old, resulting in similar SL at long muscle lengths.

**Summary Statement:** This study sought to explain the increased passive tension observed for muscles of older individuals owing to age-related changes to muscle architecture.

## Introduction

Throughout the lifespan there is a loss of muscle mass and alterations to the structural components of the neuromuscular system resulting in impaired contractile function and performance in old age (Power et al., 2013). Moreover, in old age there appears to be elevated passive tension for a given muscle length (Marcucci & Reggiani 2020). Aging muscles undergo structural remodeling, whereby muscle fascicle lengths become shorter in old as compared with young (Hooper 1981; Narici et al., 2003). A potentially underappreciated factor for increased passive tension in old age could be decreased muscle fascicle lengths owing to either a decrease in the number of sarcomeres in series and/or shorter sarcomere lengths. Thus, the fascicles and sarcomeres of muscles from old rats may experience greater relative length changes for a given displacement or joint angular rotation, resulting in increased passive tension as compared with young.

Across structural levels, passive mechanical properties of skeletal muscle are altered with aging. Investigations at the whole muscle and single fibre level have demonstrated an age-related increase in passive tension (Alnaqeeb et al., 1984; Kovanen & Suominen, 1988; Blanpied & Smidt, 1993; Gosselin et al., 1998; Brown et al., 1999; Valour and Pousson, 2003; Ochala et al., 2004; Lim et al., 2019; Noonan et al., 2020). However, others have demonstrated no age-related difference in whole muscle (Brown et al., 1999), or single muscle fibre passive tension (Wood et al., 2014). Two recent studies reported elevated passive tension in single muscle fibres in old humans (Lim et al., 2019; Noonan et al., 2020). Lim et al. (2019) reported greater passive tension and increased viscoelastic function in single muscle fibers in old compared to young. More recently, Noonan et al. (2020) reported at shorter sarcomere lengths (1.9 to 2.65 μm) single muscle fibres from old humans demonstrated a greater passive stiffness compared to young, which led to a greater passive tension in the older group at sarcomere lengths between 2.1 and 3.55 μm. Meanwhile, at longer sarcomere lengths there was no difference in the passive elastic modulus or passive tension between young and old. In contrast, Pavan et al. (2020) noted no age-related difference at the single fibre level, but passive tension of fiber bundles from old was greater compared to young across all sarcomere lengths, owing to collagen induced stiffening of the extracellular matrix. It is important to note, at the joint level, older adults have greater musculotendinous stiffness (Ochala et al., 2004; Marcucci & Reggiani 2020), and experience an earlier increase in passive force for a given joint rotation (Gajdosik et al., 2004) than that experienced by young. At the whole muscle and joint level, muscle architecture contributes relatively more to contractile function than single fibre properties (Narici et al. 2016). Thus, previously reported age-related increases in passive tension at the whole muscle level may be caused by shorter fascicles owing to reductions in the number of sarcomeres, or sarcomere length, resulting in sarcomeric structures (i.e., titin) experiencing greater stretch.

Therefore, the purpose of this study was to compare the medial gastrocnemius of young and old rats to determine differences in fascicle length, serial sarcomere numbers and the sarcomere length at which peak force is obtained (optimal reference length; RL), and record passive tension from short to long muscle lengths. It was hypothesized that fascicle length in old rats would be shorter compared with young owing to fewer sarcomeres in series, and the sarcomere length at RL would not differ between groups. Furthermore, passive tension was hypothesized to be greater in old as compared with young at long muscle lengths owing to the greater relative stretch an absolute length change will impose on shorter fascicles.

## Methods

### Animals

Barrier-reared male Fisher 344xBrown Norway (FBN) hybrid rats were obtained from the National Institutes of Aging facilities at Harlan Sprague-Dawley Inc. (Indianapolis, IN). Animals were housed at the University of Calgary in conventional housing, one per cage for less than 30 days on average before sacrifice. Housing conditions consisted of a 12 h:12 h dark–light cycle while the temperature was maintained at 22±2^°^C. Animals were provided food and water ad libitum and allowed to recover from shipment for at least 2 weeks before experimentation began. During this time, the animals were carefully observed and weighed weekly to ensure none exhibited signs of failure to thrive, such as precipitous weight loss, disinterest in the environment, or unexpected gait alterations. All procedures were approved by an animal care committee at the University of Calgary (AC13-0252).

### Surgical Preparation

9 young (8.9 ± 0.6 months; 504 ± 48g, equivalent human years ∼20 y) and 8 old (32.3 ± 0.8 months; 517 ± 50 g, equivalent human years ∼75-80 y) rats were anesthetized, the medial gastrocnemius muscle was surgically isolated and the Achilles tendon was tied in series with a force transducer for measurement of contractile properties *in situ* at 37 ± 1°C as described previously in great detail by our group: https://www.jove.com/video/3167 (MacIntosh et al., 2011).

### Electrical Stimulation Procedures

The distal stump of the sciatic nerve was stimulated with 50-μs pulses of supramaximal intensity to assure consistent activation of all motor units. The stimulating voltage was set at 1.5 × the maximum voltage (the lowest voltage that activated all motor units) to ensure maximal activation of all motor units with each stimulus. A series of double-pulse contractions (5 to 10 ms delay) at differing muscle lengths (described below) with a 20-s rest between contractions was then used to determine the muscle length that elicited the greatest active force. This length is referred to as the reference length (RL). These contractions have been shown to yield an optimal reference length, which is similar to that obtained with longer duration tetanic contractions, while preventing fatigue (MacIntosh et al., 2011).

### Passive Force Measures

Passive force was recorded at nine muscle lengths relative to the length producing optimal active force: RL − 4 mm, RL − 3 mm, RL − 2 mm, RL − 1 mm, RL, RL + 1 mm, RL + 2 mm, RL + 3 mm, RL + 4 mm. The 4 mm shortened and stretched positions used in the present study represents normal range of motion for the rat ankle joint, with the knee at 155^°^ +4 mm would represent a maximal terminal ankle joint angle in the rat of ∼26^°^ plantar flexion and −4 mm would represent ∼150^°^ of plantar flexion (Woittiez et al., 1985). Additionally, excursion of the gastrocnemius in rodents well approximates length changes of the human plantar flexors (Hu et al., 2017). To allow for dissipation of force transients owing to stress-relaxation, passive force was recorded 120 s following each of the length steps. The force transducer (Entran strain gauges (Entran Devices Inc. NJ, USA) bonded to a stainless steel lever) was mounted on a translation table that could be precisely positioned by a computer-controlled stepper motor (model MD2, Arrick Robotics Systems, Hurst, TX). The output of the force transducer was amplified (model PM-1000, CWE) and then analog-to-digital converted at 4,000 Hz (analog-to-digital board, PCI-MIO-16E-4 National Instruments, Austin, TX).

### Muscle Fixing

After all passive force and length measurements had been obtained, the animals were sacrificed and the hind limb was immediately placed in a VWR 10% Formalin (fixative) solution at the muscle length corresponding to RL. After a 1 h period of fixation, the MG muscle was dissected from the hind limb and firmly secured to a wooden applicator stick at RL and allowed to fix for 2 weeks in a VWR 10% Formalin solution. The muscles were then dissected into 4 lengthwise sections medial and lateral of the center of each MG muscle belly. After a 4 hour 30% nitric acid digestion process, 5 individual fascicles from each muscle section were isolated and placed on slides for sarcomere length measurement at 5 locations along the fascicle by laser diffraction (Lieber et al., 1984). Fascicle length measurements were taken using a digital camera and Matrox imaging software. Serial sarcomere number was calculated by dividing the fascicle length by the average sarcomere length. In total 20 fascicle length and 100 sarcomere length measurements were obtained from each muscle.

### Statistical Analyses

Unpaired two tailed t-tests were used to assess age-related differences in muscle architecture measures (muscle weight, fascicle length, sarcomere length, and serial sarcomere number). Differences in passive force between young and old across muscle length was assessed using a two-way ANOVA. For all tests, α was set to *p*<0.05.

## Results

### Muscle Architecture

Despite a similar body mass for young and old rats, the old rats had ∼34% less MG muscle mass (690 ± 40 mg) as compared with young (1040 ± 80 mg; *p*<0.05), indicating significant age-related muscle atrophy. A reduction in muscle fascicle length of ∼14% was observed in the old rats when compared to young (*p*<0.05; Figure 1 A.). There was no difference in average sarcomere lengths at RL between groups (*p*>0.05; Figure 1 B.). The MG of old rats had ∼10% fewer sarcomeres in series when compared to young rats (*p*<0.05; Figure 1 C.). Therefore, the shorter fascicle length in old muscle is driven by a reduction in serial sarcomere numbers, while sarcomere length at RL was ∼4% shorter in old, this difference was not statistically significant (Figure 1 D.).

**Figure 1.**
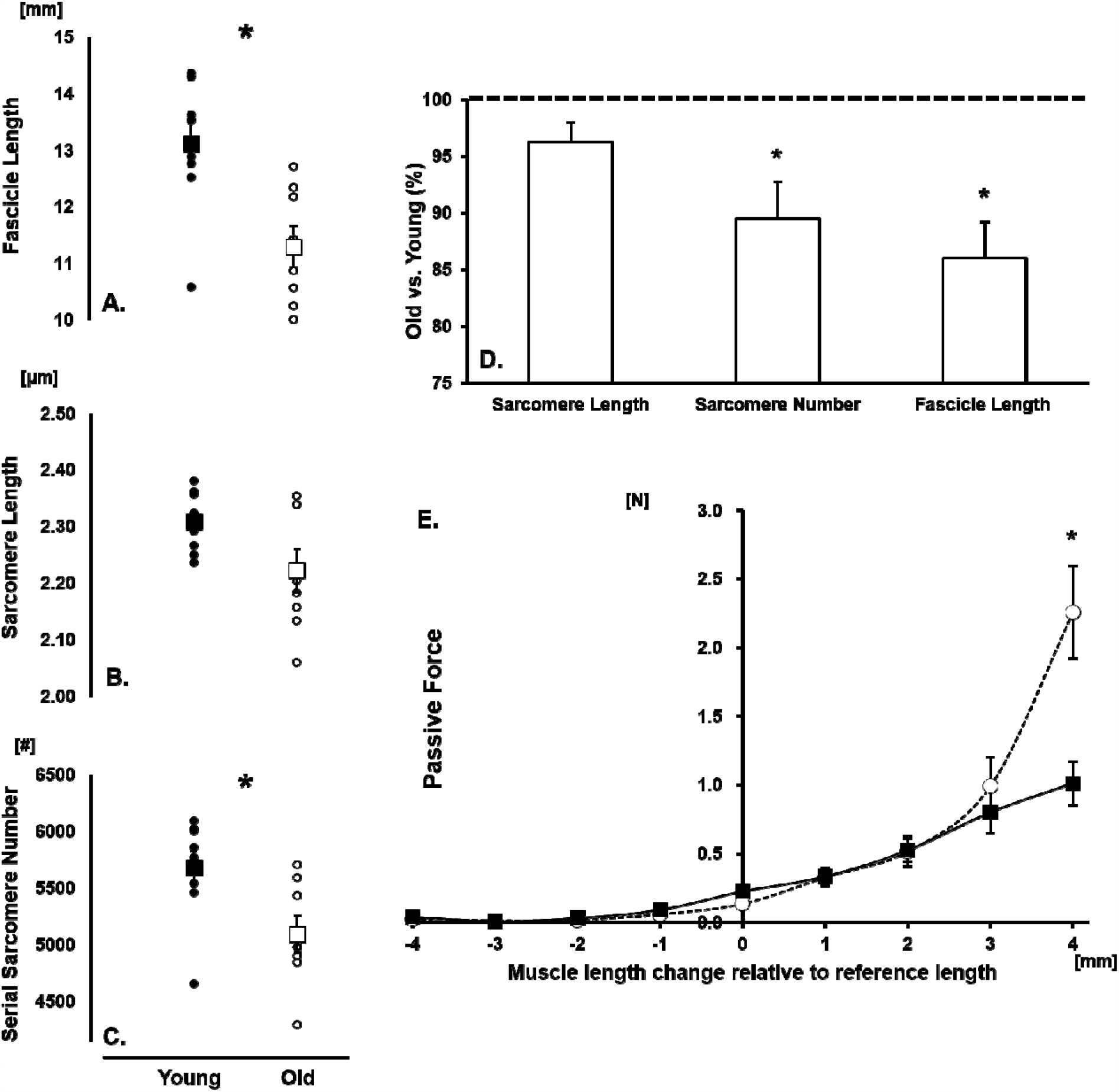
**(A.) Fascicle Length**. Fascicle length at the optimal muscle length for force production (reference length; RL) was determined using twitch doublets. The muscle was fixed and fascicle length determined by dissection, imaging and digitizing. Fascicles from Old were shorter than Young (*p<*0.05). **(B.) Sarcomere length**. Average sarcomere lengths of th dissected fascicles were measured using laser diffraction. There was no age-associated difference in sarcomere length (*p>*0.05). **(C.) Serial sarcomere number**. Sarcomere number was determined as the quotient of fascicle lengths and sarcomere lengths. Old had fewer sarcomeres in series as compared with young (*p<*0.05). **(D.)** Percent difference in sarcomere length, sarcomere number and fascicle length from the MG of Old rats relative to young (100%). **(E.) Passive force**. Passive force was determined using 1mm length steps from the reference (optimal) length. At muscle length steps beyond 3mm, Old showed significantly greater passive force as compared with Young (*p<*0.05). *Young rat data are represented as filled (black) symbols, Old rat data are represented as open (white) symbols. Significance indicated by * (p<0*.*05). All data are reported as Mean ± SEM*

### Passive Tension

There was an interaction of Age and Muscle Length for passive tension, whereby old rats experienced up to ∼125% greater passive tension at long muscle lengths (>RL+3mm) as compared with young (p<0.05; Figure 1 E.). There was no difference in passive tension at short muscle lengths between groups.

## Discussion

The present study was designed to investigate age-related alterations to muscle architecture and passive muscle tension. Using a Fisher 344xBrown Norway rat model, we show that the age-associated reduction in muscle fascicle length is primarily owing to fewer sarcomeres arranged in series, without a significant difference in resting sarcomere length. Additionally, passive force was greater in muscles from the old rats as compared with young at long muscle lengths. It was expected that the shorter fascicles and fewer serial sarcomeres in muscles from old experienced greater relative length changes for a given displacement, resulting in increased passive tension at long muscle lengths as compared with young. However, the ∼4% shorter sarcomere lengths in muscles from old, as compared with young may have dampened the effects of shorter fascicles in old minimizing any ‘overstretching’ at long lengths. For the 4 mm absolute length change, on average, fascicles from young were stretched ∼30% while old were stretched ∼36%, meaning muscles from both young and old were stretched to an average sarcomere length of ∼3.01 µm. Therefore, in the present study, differences in sarcomere length at the long muscle length are likely not the explanation for the age-related differences in passive force. This would indicate that titin, and other load bearing sarcomeric proteins, are not stretched more in old than young, and the elevated passive tension is likely collagen based. The collagen matrix likely remodeled its slack length in the old muscle, resulting in this non-contractile tissue experiencing greater strain than young when stretched to long muscle lengths. The elevated passive tension in old age, may be advantageous for joint stability and force production, thus allowing for a greater contribution of passive force to total force production compensating for muscle weakness in old age, as is often suggested for the age-related maintenance of eccentric strength phenomenon (Power et al., 2016).

### Muscle fascicle lengths are reduced in old age owing to a loss of serial sarcomeres

The reported fascicle lengths at optimal length in the present study is similar to those reported previously (13 mm) for the medial gastrocnemius in the rat (Woittiez et al., 1985). As well, the age-related reduction in fascicle length of ∼14% observed in this experiment was consistent with previously published findings of a 10-17% reduction in fascicle lengths in old humans (Narici et al., 2003; Power et al., 2013), and rats (Hooper 1981). Since sarcomeres have an optimal length for force production based on the overlap of thick and thin filaments (Burkholder & Lieber 2001), and vertebrates exhibit very little range for this optimal sarcomere length, no age-related change in resting sarcomere length was expected. Accordingly, we did not observe a significant difference in resting sarcomere length between young and old. Therefore, the ∼14% shorter fascicle lengths in old age (Figure 1 A) was expected to be primarily due to a loss of sarcomeres in series. Specifically, we observed a ∼10% reduction in the number of serial sarcomeres (Figure 1 C). Sarcomerogenesis is critical to muscle function as this process gradually re-positions the muscle, based on environmental constraints, to maintain optimal overlap of actin and myosin filaments to maximize cross bridge formation and force output (Gordon et al., 1966). An important point must be made regarding the range of sarcomere lengths on the plateau of the force length relationship. For rat skeletal muscle, assuming an actin length of 1.09 μm (Herzog et al., 1992; Ter Keurs 1984) one would presume maximal force would be obtained at sarcomere lengths between 2.28 *µ*m and 2.45 *µ*m, which corresponds to the sarcomere length observed in the present study (Figure 1 B.). Since, the plateau of the force-length curve contains a range of sarcomere lengths (2.28 – 2.45 µm), the ∼ 4% sarcomere length difference between young and old falls within that range, and likely does not affect optimal force production.

### Passive tension is elevated in the muscles of old at long lengths as compared with young

Passive tension was greater in the muscles of old as compared with young, but only at long muscle lengths. A variety of factors contribute to a muscle’s passive mechanical properties, including the structure and composition of the giant protein titin (Brynnel et al. (2019), and the collagen-based extra-cellular matrix (Prado et al., 2005; Gillies and Lieber, 2011; Meyer and Lieber, 2018). The epimysium of muscles has been reported to be stiffer in old compared to young rats (Gao et al., 2008), and similarly, Alnaqeeb et al. (1984) showed that rat extensor digitorum longus (EDL) and soleus stiffness increased with old age and was related to an increased total collagen concentration. Despite these age-related changes in connective tissue, muscle architecture is thought to play an important role in dictating mechanical properties on the whole muscle/joint level (Narici et al. 2016), especially when considered in terms of absolute excursions. Everything else being equal, the shorter fascicles in old age are stretched to a relatively greater magnitude compared to the long fascicles of the young, which, if sarcomere length remains proportional, translates into greater average SL and potentially greater passive tension at long muscle lengths. In the present study, owing to a non-significant age-related reduction in SL, this effect was buffered.

### Functional implications of fascicle length

Serial sarcomere number is important when considering the sarcomere force-length relationship (Gordon et al., 1966). For example, if the limbs of young and old were at matched joint angles such as occurs *in vivo*, assuming a given joint angular displacement produces a similar sarcomere excursion across age, the shorter fascicles in the old rats would mean that optimal length could be at the same, or smaller joint angles (i.e., shorter overall muscle length), and the sarcomeres would be stretched more, to longer lengths. Based on the sarcomere force-length relationship, this would mean at the sarcomere level, at longer muscle lengths, older adults may be operating further on the descending limb of the force-length relationship, especially during sub-maximal contractions (MacDougal et al., 2020), thus generating less active force but more passive force ultimately contributing to muscle weakness throughout a functional range of motion. Additionally, during stretch, where aged muscle sarcomeres are being pulled to relatively longer lengths than young, passive force-producing elements such as the giant protein titin may play a larger role in increased passive tension in older adults (Kellermayer et al., 1997). Moreover, it has been proposed that alterations to tendon compliance with old age may have a compensatory effect, where the shorter fascicles and fewer sarcomeres of aged muscle are stretched less, or shorten more during a fixed-end contraction (Thom et al., 2007). This age-related increase in tendon compliance in old age would assist in maintaining the sarcomere filament overlap to be closer to the optimal force production length (Thom et al., 2007).

During growth, sarcomere lengths remain relatively unchanged, assuming the thick and thin filaments reach optimal overlap (Williams & Goldspink, 1971). Further increases in fascicle length are accommodated by the addition of sarcomeres in series. In the present study, fascicle length decreased in old rats by reducing serial sarcomere numbers, while similar to growth, sarcomere length remains unchanged at maximal force capacity of the muscle. As stated previously (Hooper, 1981), this loss of serial sarcomeres may represent a period of *‘degrowth’*. The question then arises – why are muscle fascicle lengths in old muscle shorter than young? A few testable hypotheses include: *(1)* a systemic reduction in the animal’s ankle range of motion during everyday activities, thus inhibiting sarcomerogenesis (Koh & Herzog, 1998), increasing muscle stiffness, and reducing joint mobility; *(2)* a reduction in active force, minimizing tension and stretching of the fascicles during ambulation; and *(3)* a downregulation of mechanotransduction protein signalling blunting environmental stimuli for growth. These questions are beyond the scope of this short communication, but may provide some insight into sarcomerogenesis and plasticity of aging muscle.

### Optimal sarcomere length in muscle from old rats

One possible explanation for the 4% shorter sarcomeres (albeit, while not statistically significant, influenced estimates of sarcomere length when stretched to long lengths) at L_o_ in old could be related to a reduced myosin concentration (D’Antona et al., 2003.). Although speculative, if the old muscle has fewer myosin heads available to form cross-bridges, the sarcomere force length relationship could be shifted to shorter lengths to possibly increase the availability of cross-bridges and thus maintain optimal overlap, at a much smaller plateau region. As well, the shorter sarcomere length in muscle from old rats may be related to a limitation of the study where the L_o_ was measured at an optimal active contraction, but the muscles were fixed in a passive state. In this same light, the passive force was taken prior to the contraction, not estimated during the active contraction. The shorter sarcomeres in old muscle could therefore be due to factors such as the series elastic property differences in young and old muscles. Due to this limitation, it was unknown as to whether sarcomere length was altered as it entered the passive state or whether the difference was indeed due to an age-related alteration to the sarcomere structure.

## Conclusion

The results of this study were in accordance with the hypotheses that as individuals age, sarcomeres lost in series, resulting in shorter fascicles. Not surprisingly, average optimal sarcomere length did not differ between groups, but might differ therefore at different muscle lengths. The functional consequences of these age-related changes could be a reduced range of motion for older adults and less force production capacity throughout the range of motion encountered in activities of daily living. These results may also help to explain the increased passive tension observed for muscles of older individuals.

## Ethics statement

All procedures were approved by an animal care committee at the University of Calgary (AC13- 0252)

## Data accessibility

Individual values of all supporting data are available upon request.

## Author contributions

All authors contributed equally to the design of the study. G.A.P., S. C., & J. R. F., performed the experiments and analyzed data. G.A.P, wrote the first draft of the manuscript, and all authors contributed equally in editing, and approving the final version for submission.

## Acknowledgements

We would like to thank Dr. Steve Brown for comments on a previous version of this manuscript

